# PromoR and PromoS for *E. coli* promoter recognition and classification

**DOI:** 10.1101/2023.03.05.531155

**Authors:** Xinglong Wang, Jingwen Zhou

**Affiliations:** Engineering Research Center of Ministry of Education on Food Synthetic Biotechnology and School of Biotechnology, Jiangnan University, 1800 Lihu Road, Wuxi, Jiangsu 214122, China; Science Center for Future Foods, Jiangnan University, 1800 Lihu Road, Wuxi, Jiangsu 214122, China; Jiangsu Province Engineering Research Center of Food Synthetic Biotechnology, Jiangnan University, Wuxi 214122, China

**Keywords:** Promoter engineering, promoter activity, de novo design, deep learning

## Abstract

Recognition of promoters is important for novel promoter identification, which can be used for genome annotation. Classification of strong and weak promoters can be used for screening high activity promoters. This study introduced PomoR for promoter recognition and PromoS for promoter strong and weak classification. The given two network were built based on ResNet and Attention and validated by cross-validation. PromoR and PromoS displayed an accuracy of 0.887 and 0.781, respectively.

## Introduction

Promoters are DNA sequences function as controlling protein expression level by regulating gene transcription [1]. The activity of promoters affected both protein expression and secretion. It is indeed important for engineering strong promoters for overexpression of recombinant proteins or optimization of metabolic engineering in microorganisms [2–4]. Biostatistics and bioinformatics have greatly benefited for accurately and controllable designing of promoters. In this study, we taken advantages of previous studies for classification of real and fake promoters, and predicting the strength of the given promoters [5–7]. Here, we developed PromoR for promoter and PromoS for classifying strong and weak promoters.

## Results

### PromoS: classification of strong and weak promoters

PromoS was built based on ResNet [8] and Attention [9] (Fig. 1A) for strong and weak promoter classification. PromoS was trained used RegulonDB (3382 samples) dataset including 1591 strong and 1792 weak promoters [10]. Promoters in the training set was adjusted to 50 bp to suite the length of generated promoter (Fig. 1B). Several optimization methods were attempted for model architecting and training. In short, sequential feature extraction using one hot encoding strategy outperformed using pseudo-dinucleotide composition (pseDNC) method or the combined of the two methods. Network architecture based on ResNet and attention achieved better accuracy than simply using CNN or ResNet (Fig. 1C). Hyperparameter optimization showed that the batch size of 32 and learning rate of 0.0001 gave the best result (Fig. 1C). PromoS achieved state-of-art accuracy of 0.781 (Fig. 1C).

**Figure 1.**
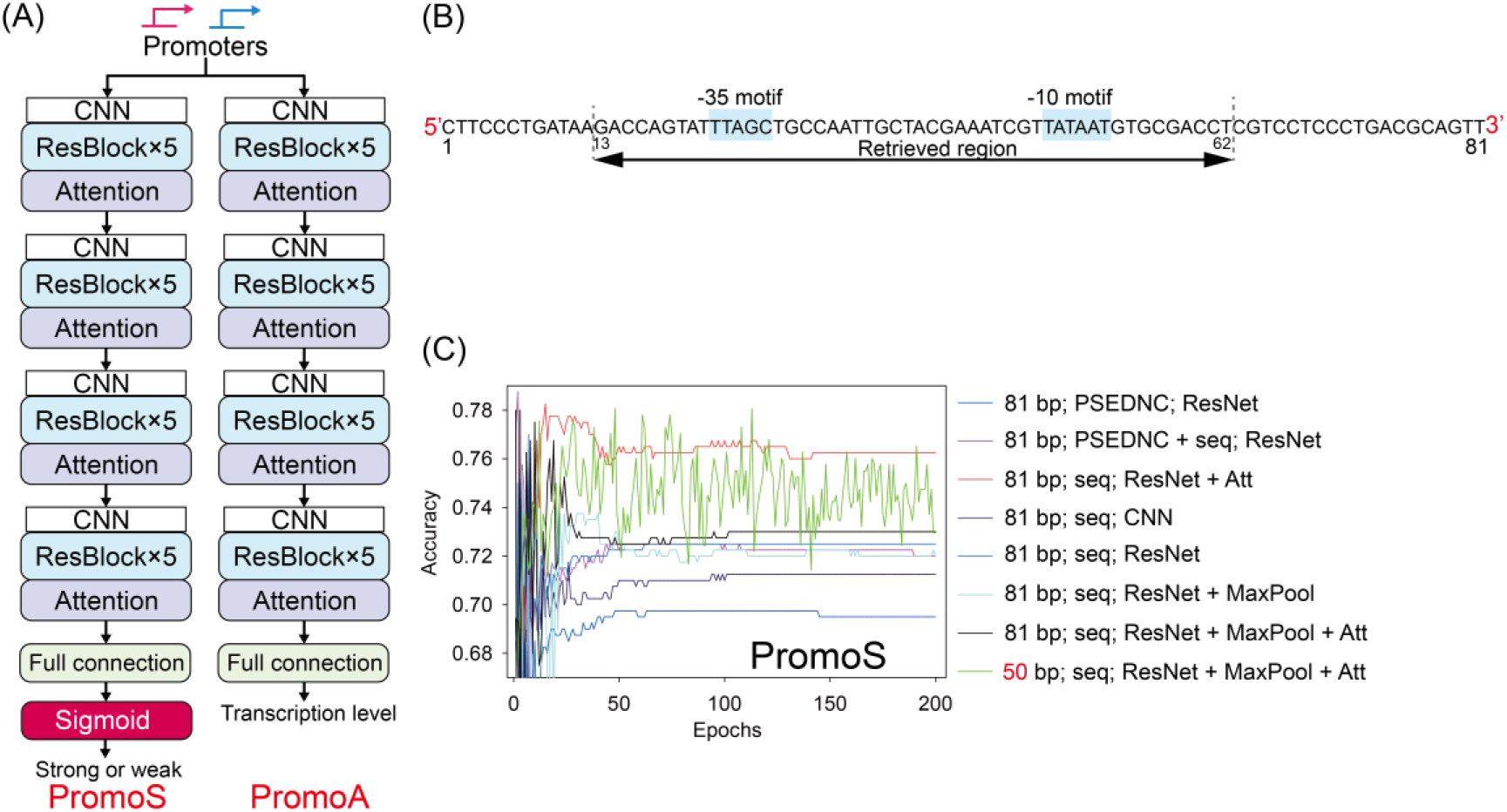
Architecting of PromoS and PromoA. (A) The network architecture of PromoS and PromoA; (B) Retrieving 50 bp from the original promoter sequence based on −35 and −10 motif distribution; (C) Attempted methods for PromoS optimization.

### PromoR: classification of real and fake promoters

Recognition of real and fake promoters can critically evaluate the quality of generated promoters. PromoR was built for classifying real and fake promoters, the network taking advantage of PromoS with minor differences (Fig. 1A). This network was initially trained using 3382 promoters and 3382 non-promoters from RegulonDB [10], but adjusted the promoter length to 50 bp regarding to the −35 and −10 motifs distribution in natural promoters (Fig 1B). PromoR achieved a state-of-art accuracy of 0.853 which is comparable to previous reported highest values.

PromoR was validated using an independent dataset organized by mixing 11,884 natural promoters [11] and 11,884 non-promoters generated by randomly retrieving 50 length sequence from the non-promoter (81 bp) in RegulonDB. The validation set was non-reductant itself and shared non-overlaps with training set. PromoR displayed an accuracy of 0.767, which was much lower than the cross-validation results. The failed samples showed similar values of 11.4% and 11.9% for real and fake promoters, highlighted weaker performance of PromoR towards more diversified samples. To reinforce the capability of PromoR, the validation and training set were combined for retraining PromoR, and randomly split 20% of data for cross-validation. Retrained PromoR with enlarged dataset has dramatically promoted its accuracy, which reach to 0.887. These results in agree to previous studies demonstrating training model with exclusive data can enhance its performance [12].

## Conclusion

In the present study, we introduced PromoR and PromoS for promoter recognition and strength prediction. The network and training set were optimized in order to achieve high accuracy. Our result showed PromoR and PromoS displayed an accuracy of 0.887 and 0.781, respectively. The two given network may further benefit *E. coli* genome annotation and screening high activity promoters.

## Acknowledgements

This study was funded by the National Key Research and Development Program of China (2019YFA0904900) and the Starry Night Science Fund of Zhejiang University Shanghai Institute for Advanced Study (Grant No. SN-ZJU-SIAS-0013).

## Author information

### Authors and Affiliations

**Science Center for Future Foods, Jiangnan University, 1800 Lihu Road, Wuxi, Jiangsu 214122, China**

Xinglong Wang, Jingwen Zhou

**School of biotechnology, Jiangnan University, 1800 Lihu Road, Wuxi, Jiangsu 214122, China**

Xinglong Wang, Jingwen Zhou

**Jiangsu Province Engineering Research Center of Food Synthetic Biotechnology, Jiangnan University, Wuxi 214122, China**

Jingwen Zhou

**Corresponding author**

Correspondence to Xinglong Wang and Jingwen Zhou

## Ethics declarations

### Availability of data and material

All data related to this study has been included in the manuscript.

## References

1. Keasling JD (1999) Gene-expression tools for the metabolic engineering of bacteria. Trends in Biotechnology 17: 452–460.

2. Jin L-Q, Jin W-R, Ma Z-C, Shen Q, Cai X, et al (2019) Promoter engineering strategies for the overproduction of valuable metabolites in microbes. Applied Microbiology and Biotechnology 103: 8725–8736.

3. Keasling JD, García Martín H, Lee TS, Mukhopadhyay A, Singer SW, et al (2021) Microbial production of advanced biofuels. Nature reviews Microbiology

4. Even D, Kedmi A, Basch-Barzilay S, Ideses D, Tikotzki R, et al (2016) Engineered Promoters for Potent Transient Overexpression. PLoS ONE 11

5. Chen W, Feng P, Lin H, Chou K-C (2013) iRSpot-PseDNC: identify recombination spots with pseudo dinucleotide composition. Nucleic Acids Research 41: e68–e68.

6. Zhang Z-m, Zhao J-p, Wei P-J, Zheng C-H (2022) iPromoter-CLA: Identifying promoters and their strength by deep capsule networks with bidirectional long short-term memory. Computer Methods and Programs in Biomedicine 226: 107087.

7. Tayara H, Tahir M, Chong KT (2020) Identification of prokaryotic promoters and their strength by integrating heterogeneous features. Genomics 112: 1396–1403.

8. He K, Zhang X, Ren S, Sun J (2015) Deep Residual Learning for Image Recognition. 2016 IEEE Conference on Computer Vision and Pattern Recognition (CVPR): 770–778.

9. Vaswani A, Shazeer NM, Parmar N, Uszkoreit J, Jones L, et al (2017) Attention is All you Need. ArXiv abs/1706.03762

10. Gama-Castro S, Salgado H, Santos-Zavaleta A, Ledezma-Tejeida D, Muñiz-Rascado L, et al (2015) RegulonDB version 9.0: high-level integration of gene regulation, coexpression, motif clustering and beyond. Nucleic Acids Research 44: D133–D143.

11. Thomason MK, Bischler T, Eisenbart SK, Förstner KU, Zhang A, et al (2014) Global Transcriptional Start Site Mapping Using Differential RNA Sequencing Reveals Novel Antisense RNAs in Escherichia coli. Journal of Bacteriology 197: 18–28.

12. Ng W, Minasny B, Mendes WdS, Demattê JAM (2020) The influence of training sample size on the accuracy of deep learning models for the prediction of soil properties with near-infrared spectroscopy data.

